# Genome scale metabolic modelling of human gut microbes to inform rational community design

**DOI:** 10.1101/2024.05.28.596116

**Authors:** Juan Pablo Molina Ortiz, Dale David McClure, Andrew Holmes, Scott Alan Rice, Mark Norman Read, Erin Rose Shanahan

## Abstract

The human gut microbiome impacts host health through metabolite production, notably short-chain fatty acids (SCFAs) derived from digestion-resistant carbohydrates (DRCs). While DRC supplementation offers a means to modulate the microbiome therapeutically, its effectiveness is often limited by the microbial community’s complexity and individual variability in microbiome functionality. We utilized genome-scale metabolic models (GEMs) from the AGORA collection to provide a system-level overview of the metabolic capabilities of human gut microbes in terms of carbohydrate trophic networks and propose improved therapeutic interventions, based on microbial community design.

Our study inferred the capability of AGORA strains to consume carbohydrates of varying structural complexities—including DRCs—and to produce metabolites amenable to cross-feeding, such as SCFAs. The resulting functional database indicated that DRC-degrading abilities are rare among gut microbes, suggesting that the presence or absence of specific taxa can determine the success of DRC-based interventions. Additionally, we found that metabolite production profiles exceed family-level variation, highlighting the limitations in predicting intervention outcomes based on gut microbial composition assessed at higher taxonomic levels.

In response to these findings, we integrate reverse ecology principles, network analysis and GEM community modelling to guide the design of minimal yet resilient microbial communities to better guarantee intervention response (purpose-based communities). As a proof of principle, we predicted a purpose-based community designed to enhance butyrate production when used in conjunction with DRC supplementation, that displays resilience under nutritional stress, such as amino acid restriction.

We further seeded the identified purpose-based community into modelled human microbiomes previously demonstrated to accurately predict SCFA production profiles. The analysis confirmed that such intervention significantly promotes butyrate production across samples, with those that presented a comparatively lower butyrate production pre-intervention displaying the largest increase in butyrate production after seeding. Our work highlights the potential of combining GEMs with community design to infer effective microbiome interventions, ultimately leading to improved health outcomes.

## Introduction

The human gut microbiome plays a pivotal role in health and disease (1–3), including through the production of metabolites with biological effects on the host. Evidence consistently suggests that this microbial community is strongly influenced by host diet, particularly through the intake of digestion-resistant carbohydrates (DRCs) (4–11). This offers the opportunity of modulating the microbial community to alter host health outcomes, such as via DRC supplementation, or with microbes that ferment these substrates (12–15). However, the effectiveness of such interventions is hindered by the inherent complexity of microbial communities and the variable functional capacity of microbiomes across individuals (7, 16–19).

While further efforts are needed to fully gasp human gut microbiome dynamics, and therefore successful microbiome modulations, broad principles have been established. For example, evidence supports the existence of trophic networks in this microbial community, where specific commensals can degrade DRCs (primary degraders), enabling other microbes access to less complex carbohydrates (secondary degraders) (20–22). It is also understood that gut commensals can adapt their metabolism to changes in substrate availability, and thus a microbe’s role as a primary or secondary degrader may change accordingly (9). Moreover, characteristics common to the gut environment, including both high rates of substrate availability and intense competition for resources, favour partial degradation of nutrients and accumulation of intermediate metabolites, such as formate, which can then be cross-fed to alternative populations in the community (23, 24) or made available to the host. Short chain fatty acids (SCFAs), some of the most abundant metabolites produced by human gut microbes, mostly through DRC fermentation, are associated with health promoting effects (4, 25–27). Meanwhile, other metabolites such as hydrogen sulphide (H2S), are generally considered detrimental to health (8–11). These principles underscore the need for a systems-level understanding of how nutritional inputs impact metabolic outputs in the human gut microbiome for the design of successful interventions.

Despite the popularity of probiotics, currently available formulations often yield inconsistent and limited results (28–30). This inconsistency is largely due to the complex and competitive nature of the gut microbial ecosystem, where introduced strains struggle to establish themselves and exert significant functional impact (28, 29). Single strains may also lack the necessary metabolic breadth to degrade diverse DRCs or produce sufficient beneficial metabolites, such as SCFAs. Therefore, there is a growing recognition that effective interventions may require the introduction of a tailored community of microbes that can work synergistically to degrade DRCs and produce health-promoting metabolites (31, 32).

To achieve this comprehensive view and to overcome the challenges of studying this complex ecosystem *in vivo*, researchers have increasingly turned to computational approaches. Genome-scale models (GEMs) have gained momentum in the field to address such challenges (11, 31, 33–35). GEMs are computational integrative platforms that allow the amalgamation of different data types (e.g. omics and empirical) to build and refine the metabolic network encoded in genomes. In doing so, GEMs reconcile genetic traits at the cellular level, offering insight into what is metabolically feasible for a given strain (36–38). GEMs can be modelled in isolation or as a consortium, in nutritional conditions that mimic those encountered in the human gut, while tracing intracellular fluxes and intercellular metabolic exchanges, providing mechanistic insights that would be complicated to attain through alternative methods. The resulting predictions also have the potential to inform the design of minimal, resilient microbial consortia (46–48), and optimal concurrent dietary supplementation, for microbiome-targeted intervention strategies (purpose-based communities). Well-curated GEMs are of particular importance in performing such modelling as they better capture species-specific functional capabilities and potential metabolic interactions. In this regard, AGORA, a collection of 818 highly curated GEMs of cultured and un-cultured gut microbes, has been widely applied to study the human gut microbial community and has been validated against human data (31, 39–42).

Leveraging GEMs, this study, previously made available as a preprint (43), first aims to provide a system-level overview of the metabolic capabilities of human gut microbes in terms of nutritional inputs and metabolic outputs in carbohydrate trophic networks. To do this, we comprehensively characterised the metabolic capabilities of gut commensal AGORA GEMs to a) consume carbohydrates from various levels of structural complexity, including DRCs; and b) produce metabolites that can be cross-fed, notably SCFAs. We also aimed to demonstrate the potential of GEMs for the design of a purpose-based community that is optimised for SCFA production in the context of the human gut microbial community. We show that a resulting proof of principle community is predicted to be resilient to stressors such as amino acid limitation, and enhance butyrate production in modelled human microbiomes (31). Finally, flux inspection highlights functional characteristics in modelled microbiomes with potential implications for intervention response. Our work highlights the potential of GEMs for understanding the metabolic potential of microbial communities, and leveraging this knowledge to design and assess microbial microbiome interventions in the context of the human gut microbiome.

## Methods

### Computational resources

The GEMs employed for this project were obtained from www.vmh.life (44), and are part of the *without mucins* version of AGORA (Assembly of Gut Organisms through Reconstruction and Analysis) version 1.03 (39). This was selected based on our aim of focusing on dietary-derived molecules (with the 1423 *O*-glycans represented in the standard (*with mucins*) version being out of the scope of this body of work). GEMs of strains *Clostridium sporogenes* ATCC 15579 and *Lactobacillus helveticus* DPC 4571 were excluded from this analysis due to inconsistencies, as outlined previously (42).

Metabolite flux for each GEM was modelled using Flux Balance Analysis (FBA) and the constraint-based reconstruction and analysis (COBRA) toolbox for the Python coding language COBRApy (45). FBA is a mathematical optimisation approach that maximises an objective function based on upper and lower bounds imposed on reaction fluxes (constraints). In FBA, multiple flux distributions (solutions) can achieve the same optimal objective value. We applied FBA to optimise growth in terms of biomass generation in grams of dry weight per hour (gDW/h) based on nutrient fluxes (equivalent to specific growth rate (µ) when the biomass flux is normalised to 1 gDW of biomass). COBRApy further allows a) inspection of individual metabolic reactions and metabolite consumption and synthesis and b) seamless addition or removal of specific molecules to growth media.

Community modelling was performed using MICOM (46), a python package for modelling individual GEMs in a community context. Fluxes were derived from utilising the cooperative_tradeoff method, which optimises the community metabolic network for maximum community and individual member growth. Parsimonious FBA (pFBA) was used to model community fluxes, a variation of traditional FBA, which aims to minimise the total sum of predicted fluxes that contribute to an optimal objective function, reducing the solution space by selecting the least-costly metabolic route, which is highly desirable during community modelling. MICOM cooperative_tradeoff simulations were performed with a tradeoff value of 0.99, to further constrain the solution space. COBRApy and MICOM simulations were performed with an academic license for the Gurobi solver (47) on a personal computer.

### Nutrient utilisation inference

To determine which nutrients each strain can utilize, we developed a COBRApy-based script that systematically tests the ability of GEMs to grow on specific nutrients and produce metabolites. Previously, we introduced GEMNAST (GEMs Metabolic and Nutritional Assessment), a Python-based algorithm that samples metabolic capabilities of a predetermined set of GEMs qualitatively, based on combinatorial media modifications (42). GEMNAST (https://github.com/jmol0917/GEMNAST_0.2) was devised as a high-throughput pipeline to identify auxotrophies and prototrophies in GEMs. For this study, GEMNAST was extended to assess nutrient utilisation and metabolite export capabilities on individual GEMs.

### Nutrient utilisation capabilities

A COBRApy-based python script was developed to identify molecules individual GEMs can utilise as nutrients (Figure S1 in Supplementary file 1), where each GEM is simulated in base media enriched with the potential nutrient, and constrained to utilise the added molecule, if possible. The main output of this pipeline is a Boolean table with strains as rows and test molecules as columns. If the model produces biomass above a predetermined threshold and consumes the molecule, confirmed by flux inspection, a positive outcome is recorded (1). If both conditions are not met, a negative outcome is assigned instead (0).

### Metabolite export capabilities following nutrient utilisation

We developed a pipeline to identify which metabolites can be exported by GEMs when utilizing specific nutrients, helping to infer cross-feeding potential among microbes in a community. We designed a script that enables a predetermined set of compounds to be assessed (Figure S2, Supplementary file 1). In this pipeline, each GEM is simulated in base media enriched with the potential nutrient, and constrained to i) utilise the added molecule and ii) to export (after producing) the metabolite in question, if possible. If these two conditions are fulfilled (confirmed by flux inspection) and the model produced biomass above a determined threshold, a positive outcome (1) is recorded in a Boolean table where the represented strains are recorded in rows and combinations of one molecule and one metabolite in columns. If every condition is not met, a negative outcome is assigned instead (0). It is important to note that this analysis does not trace the specific metabolic pathways leading from nutrient uptake to metabolite export. Therefore, while it confirms the capability to produce a metabolite during nutrient utilization, it does not guarantee that the metabolite is directly derived from that nutrient.

### Growth media for modelling

The previously reported Universally Defined Media (UDM, Table S1) (42), which was designed to guarantee growth of every GEM in AGORA, was modified for the purposes of this study. To accurately assess carbohydrate utilization capabilities, we modified the UDM by removing potential confounders (e.g. simple sugars) that could serve as alternative carbon sources, resulting in a carbohydrate-free UDM (cfUDM, Table S1).

### Meaningful growth

To determine a strain’s capacity to grow in media an extremely conservative threshold of 0.09 h^−1^ growth rate (eight hours doubling time) was established on the basis that any microbe growing at a lower rate would present relatively low probabilities of surviving in the human gut, where transit times average just over 24 hours, but can be shorter than 14 hours in some individuals (48). This threshold represents a theoretical minimum growth rate for gut survival as less than two replication cycles would not guarantee permanence in the human colon (49).

### Carbohydrate utilisation and metabolite export experimental design

We selected 55 carbohydrates relevant to DRCs based on the VMH database (44), where AGORA GEMs are housed (39), and literature reports. These were categorized by structural complexity into monosaccharides, oligosaccharides, and polysaccharides. Additionally, 11 metabolites were identified from the same sources for the second part of this analysis, focused on a subset of metabolites particularly relevant to fermentation of microbial-accessible carbohydrates and production of short-chain fatty acids (50). The names of these 66 molecules can be found in Table S2.

Following our customised GEMNAST script, we sampled the carbohydrate utilisation capabilities of 816 strains represented in AGORA (input a) based on the 55 carbohydrates (input c) and cfUDM as base media (input b). Each molecule was individually added to cfUDM, and molecule utilisation and growth were assessed (Figure S3). Results from this analysis are shown in Supplementary file 2.1, 2.2 and 2.3 for monosaccharides, oligosaccharides and polysaccharides, respectively.

The GEMNAST script we developed to assess metabolite export was applied to identify which of the 11 selected metabolites (input d) AGORA strains (input a) can export while utilising each monosaccharide from Table S2 (input c), with cfUDM (input b) as base media (Figure S4). Results of this part of our analysis can be found in Supplementary file 2.4.

### Classification of AGORA strains into topo-physiological categories

AGORA strains were divided into groupings (Supplementary file 2.5) based on their relationship with the host (pathogens, pathobionts) and their more prevalent anatomical site (upper GIT, large bowel, skin). This process was done manually with reference to the literature and relevant databases (51–54).

### Phylogeny assignment

Taxonomy IDs of 590 large bowel AGORA strains were retrieved from the Virtual Metabolic Human database (44) and were used to assemble a .phy file using NCBI’s online taxonomy browser (55) (Supplementary file 5) to retrieve updated phylogenetic details for such strains and families relevant for our analyses.

### t-Distributed stochastic neighbour embedding (t-SNE) and Kernel density estimation (KDE)

We used t-SNE for dimensionality reduction to visualize high-dimensional metabolic profiles, and KDE to estimate strain density distributions within the t-SNE plots. t-SNE and KDE were applied to the Boolean tables that resulted from the analysis of each category of molecules to visualise how strains from prevalent large bowel families are distributed in the space, based on their individual carbohydrate degradation/metabolite production profiles. The TSNE scikit-learn Python module was used for this purpose. We utilised the Jaccard distance and exact t-SNE, which optimises the Kullback-Leibler divergence of distributions between original and embedded spaces, to measure the dissimilarity between datapoints (56). KDE, from the SciPy, was used to estimate the number of overlapping strains in on a t-SNE plot (57).

### Heatmaps, Boxplots and Networks for results visualisation

Seaborn (58), a Python library for data visualization, based on matplotlib (59), was used to generate heatmaps and boxplots derived from the carbohydrate utilisation and metabolite production datasets and correlation matrices in the purpose-based community. Matplotlib was used to generate stacked bar charts (59). Cytoscape (60) was used to generate interactions between members of the purpose-based community.

### Flux network analysis

To identify strains in the 151-strain community that are more likely to establish a resilient purpose-based community we performed a network analysis based on predicted nutritional exchanges from community modelling. Every modelled molecule was included in this analysis. First, we calculated total in- and out-degree edges (predicted imported or exported metabolites) for every strain in this community. We then utilised this metric to identify a) hub strains, those with a high imports and exports sum (> 6.7, fourth decile); and b) strains with a balanced export:import ratio (exports/imports = 0.9-1.1), to prevent the inclusion of “cheaters” i.e. organisms that are low exporters.

### Percent change in butyrate production rates

To report the difference in butyrate production rates between two modelled communities or different nutritional contexts we rely on Equation 1 to calculate percent changes.

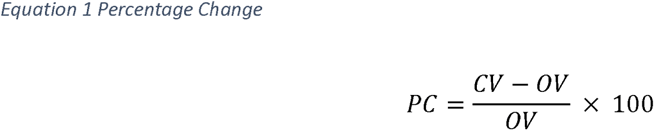

Where PC is the percent change, OV is the original value; in this case, the predicted rate for butyrate production in our purpose-based 6-strain community in cfUDM supplemented with inulin. CV is the compared value, which corresponds to alternative butyrate production rates. Absolute predicted values can be found in the corresponding sections of Supplementary file 2 as part of community modelling fluxes.

### Seeding a purpose-based GEMs community in predefined species-level human gut microbiome-derived MCMMs

We reproduced the microbial community-scale metabolic models (MCMMs) derived from human participant faecal samples from Quinn-Bohmann *et al*., Study A to D (31), based on the publicly available data and code associated with the study. Briefly, species (study A, C and D) and genus-level (Study B) taxonomy and abundances, inferred from 16S rRNA gene amplicon sequencing, were utilised to build MCMMs. When repeat data was available (studies A and B), it was modelled as independent MCMMs. This resulted in 53 original MCMMs (study A (*n* = 6), B (*n* = 29), C (*n* = 9) and D (*n* = 9)), which were modelled in standard European diet (44) supplemented with 10 mmol/gDW/h of inulin (fibre intervention). Our purpose-based community (strain-level GEMs) was then seeded into the original MCMMs at an overall abundance of 5% and then modelled in the inulin supplemented European diet (fibre + module intervention). This process was performed utilising MICOM’s multi-sample *build* and *grow* workflows with a tradeoff parameter of 0.7, derived from MICOM’s cooperative tradeoff analysis (46), which recommends the use of the highest parameter value that allows 90% or more of the taxa in a MCMM to grow. Simulated inulin supplementation and purpose-based community relative abundance seeding values are the same utilised by Quinn-Bohmann *et al*. Fluxes that corresponded to different samples from the same participant were then averaged for later analysis. This resulted in a final total of 30 MCMMs (studies A (*n* = 2), B (*n* = 10), C (*n* = 9) and D (*n* = 9)).

A tradeoff parameter of 0.7 constrains MCMMs to grow at a 70% maximum community growth rate, which can translate into a higher number of individual GEMs simulating non-zero growth rates. However, this also translates into a larger solution space, where metabolite production rates can vary widely. To address this, we further simulated MCMMs with butyrate production as the objective function, while constraining individual GEM growth to the growth rates obtained in the unconstrained *grow* workflow, which predicts the theoretical maximum butyrate-production a given MCMM can achieve. Resulting growth rates and predicted fluxes were then divided by a factor of 10 to account for faecal microenvironment dilution (31), while preventing numerical instability during modelling, as reported previously (11).

## Results

### *In silico* culturing with a stratified carbohydrate dimension to characterize metabolic capabilities

DRCs, commonly termed dietary fibre, are not digested in the human small intestine and therefore pass into the large bowel. Not all DRCs can be utilised by gut microbes, but those that can be metabolised (microbiota-accessible carbohydrates) represent key carbon and energy sources for microbes in the large bowel. Comprehensive *in silico* characterisation of the functional traits of AGORA strains in the carbohydrate nutritional dimension requires equally comprehensive representation of this group of molecules. To adequately represent the structural complexity within carbohydrates, relevant to the human gut microbiota, we designated three categories based on their chemical structure. Each of these categories was populated with compounds from a list of nutrient requirements used for the curation of AGORA GEMs (44), totalling 55 compounds (Table S2, Supplementary file 1):

- Polysaccharides (25 compounds): carbohydrates with over ten monosaccharides in their structure.
- Oligosaccharides (14 compounds): molecules composed of two to ten monosaccharides.
- Monosaccharides (16 compounds): single sugar units that cannot be further broken down by hydrolysis.

To characterise carbohydrate utilisation by AGORA strains, we extended our GEMNAST pipeline (42), which facilitates inference of metabolic traits encoded in GEMs (Methods). To infer if a strain is capable of utilising a given carbohydrate, its GEM counterpart should predict that the strain can ‘grow’ (produce biomass) above a predetermined threshold while utilising the molecule. Using this pipeline, each AGORA strain was modelled for ‘growth’ in each of the 55 selected carbohydrates, totalling 53,040 *in silico* cultures (Supplementary files 2.1 to 2.3).

Genome-scale metabolic modelling also allows inference of metabolites, products of microbial metabolism that can accumulate in the environment from nutrient foraging. Leveraging this functionality, we aimed to qualitatively infer intermediate and terminal metabolites that can be produced and exported when large bowel strains utilise specific monosaccharides, due to their importance as cross-feeding currency and means of interaction with the host. The literature suggests that both terminal metabolites (e.g. SCFAs) and even intermediate metabolites can drive metabolic interactions in the human gut (24) and other microbial systems (61). To obtain a more comprehensive trophic network perspective, and given that different metabolic strategies may lead to the production of alternative metabolites (18, 24, 62), we relied on modelling constraints to ‘induce’ individual GEMs to export a given metabolite while utilising a monosaccharide, if metabolically feasible. This provides insight into whether it is *possible* for a strain’s metabolic network to adopt such a metabolic phenotype in the given nutrient conditions. A total of 11 metabolites (four intermediate and seven terminal, Table S2) were selected for this analysis based on AGORA curation data and literature reports (39), focused on a subset of metabolites particularly relevant to fermentation of microbial-accessible carbohydrates and production of short-chain fatty acids (50). This resulted in 105,248 *in silico* cultures (Supplementary file 2.4).

The focus of our analysis is the human large intestinal microbial community, as this is a key target for interventions. Although AGORA comprises GEMs of strains that have been identified in, and/or can inhabit the human large intestine, we focused our analysis on taxa that could be considered stable colonisers of the large bowel, excluding for example, oral taxa and known pathogens (Supplementary file 2.5). Comparison of the carbohydrate utilisation capabilities between large bowel strains and other groups of strains from AGORA can be found in Table S3 and Figure S5.

### Primary degraders as determinants of intervention success

We analysed the distribution of carbohydrate utilisation capabilities in strains from the large bowel to better understand their predisposition to different ecological roles. At a broad level we found that carbohydrate utilisation capabilities are increasingly rare at higher levels of nutritional complexity. For large bowel strains, 173 out of 598 (28.93%) could utilise one or more polysaccharides, a proportion that progressively increased for less complex groups of molecules (Figure 1A-C). This suggests that a larger portion of large bowel microbiota are predisposed to occupy secondary degrader roles, and that fewer strains possess the functional traits needed for a primary degrader role.

**Figure 1.**
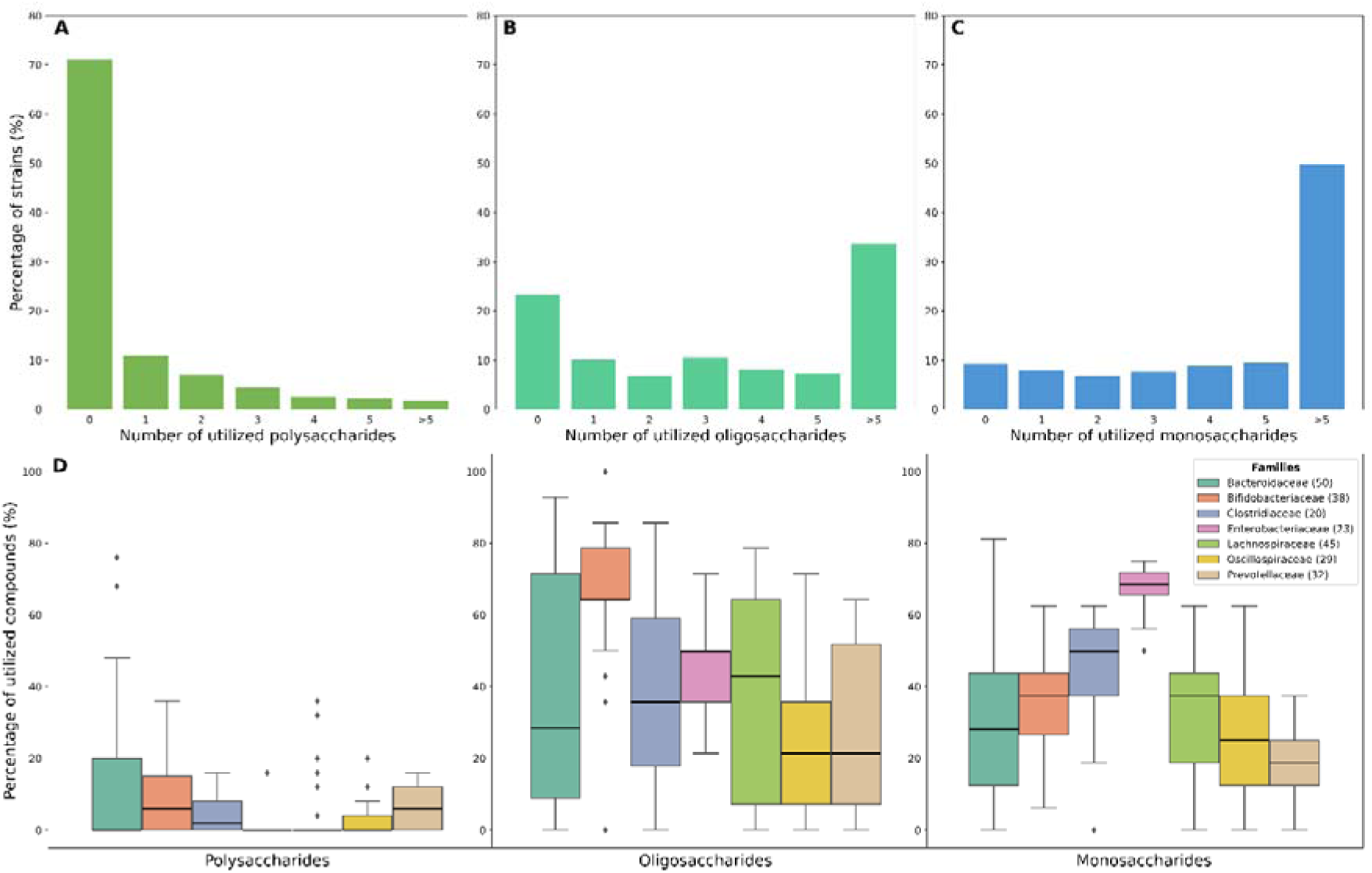
Carbohydrate utilisation capabilites of stable large bowel colonisers in AGORA. A) Number of strains capable of utilising at least one carbohydrate from the given structural category. A lower number of strains are inferred to utilise molecules within a group as structural complexity increases. A-C) Percentage of strains inferred to utilise a given number of carbohydrates within molecular structure categories. A lower number of strains are inferred to utilise molecules within a group as structural complexity increases based on the percentage of strains that utilise zero compounds in that group of molecules. D) Boxplot showing average proportion of assessed nutrients from each structural category that can be utilised by strains within each bacterial family. Family median is shown in bold.

Of the strains that could utilise polysaccharides (*n* = 173) about half can only consume one of the 25 polysaccharides we analysed (Figure 1A). Panels B and C in Figure 1 show that the opposite occurs in less complex carbohydrate dimensions, where most of the strains can consume more than five nutrients in each category. When focusing on bacterial families that are both prevalent in the human gut and in AGORA (Figure 1D), we find a similar trend, as the median percentage of polysaccharides that can be used by each strain within these families is low or zero. The presence of outliers suggests that polysaccharide degradation capabilities are the exception, not the rule, among common gut commensals. Notably, we evidenced strain-level differences in polysaccharide utilisation capabilities in, for example, *Eubacterium rectale*, *Clostridium symbiosum* and *Bifidobacterium breve* strains. The opposite occurs in less complex dimensions, where strains can consistently consume from 20% to over 60% of carbohydrates in the oligo- and monosaccharide groups across each of the bacterial families.

The implication of these findings for nutrition and DRC-based interventions is that the successful incorporation of a polysaccharide (and its derivatives) into trophic networks might depend on very select microbial strains. If such strains are missing from a given individual’s microbiome, a given DRC is less likely to result in modulation of the microbiota or downstream health outcomes. On the other hand, once a supplemented DRC has been broken into its less complex constituents it is likely that such derivatives will be accessible to a wider portion of community members.

### Metabolite production profiles in the human gut exceed AGORA phylogeny

A second part of our analysis focuses on the fermentation/SCFA related metabolic profiles large bowel strains can produce when utilising the range of monosaccharides considered in our assessment. Our results showed that the majority of large bowel strains (542 of 598 tested) can adopt metabolic states that allow them to grow above a minimum threshold while producing and exporting at least one of the 11 metabolites. Of these strains, the vast majority (*n* = 530) presented a qualitative metabolite production profile that remained consistent irrespective of which monosaccharide was available in the medium (Supplementary file 2.6). The production profile represents a set of viable metabolic options each strain may use, noting that not all metabolites will be produced simultaneously. However, we identified 129 unique sets of metabolite combinations in total, far exceeding family level variation represented in AGORA strains (39). Many metabolic profiles were unique to a particular strain, and only 29 metabolite production profiles were shared among more than four strains. The largest groups of strains sharing the same metabolite profile ranged from 48 to 20 members (Figure 2A). Notably, most of these groups are composed of members of the same family. Overall, our analysis predicts that on average 316 strains can produce each of the tested metabolites, however only 82 of the large bowel strains tested can produce butyrate, considered a key SCFA for intestinal health (63–65), in the tested environment (Figure S6 in Supplementary file 1).

**Figure 2.**
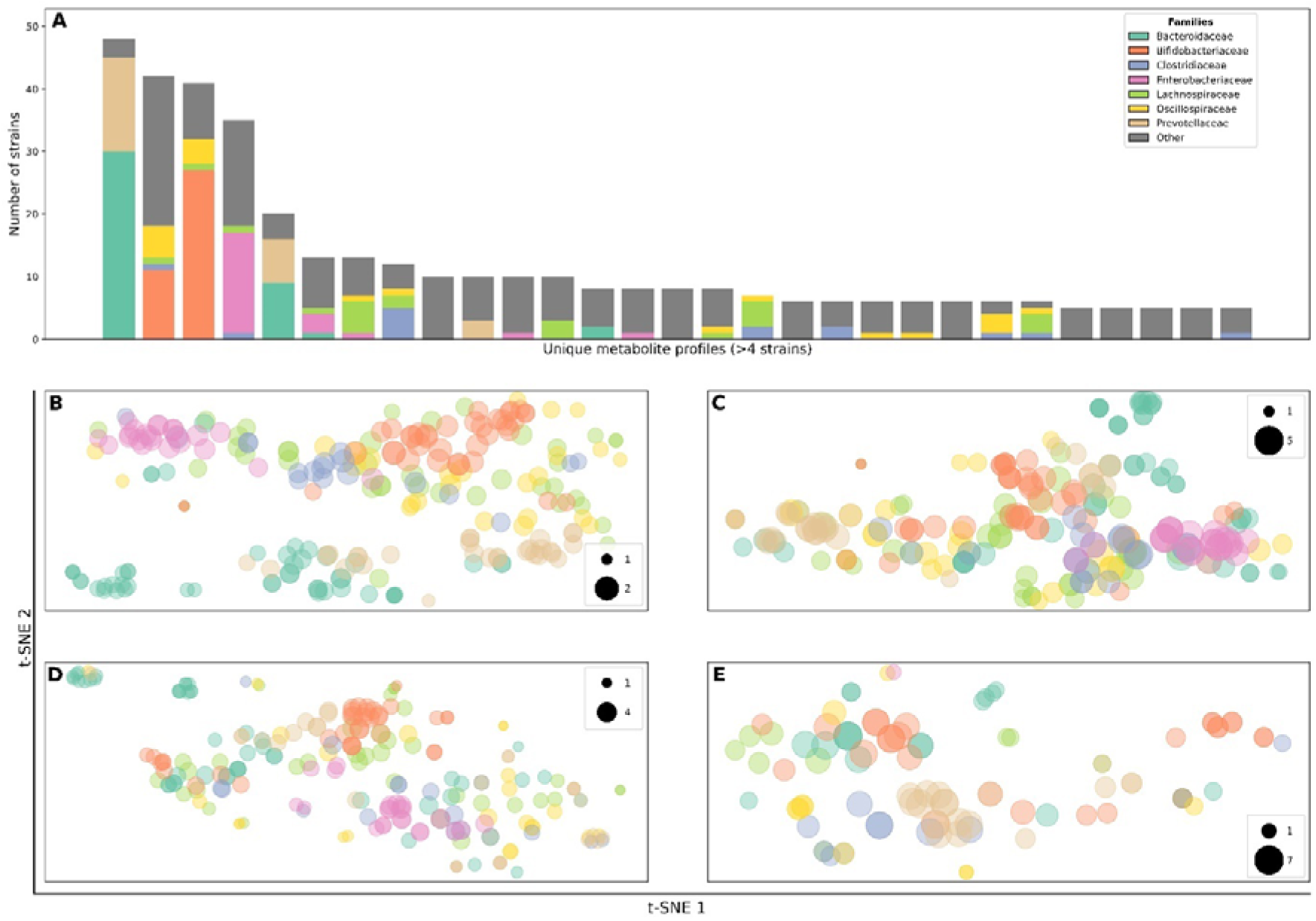
Carbohydrate utilisation profiles among prominent human gut microbiome families represented in AGORA. A) Metabolite production profiles (sets) shared by more than four GEMs. Number of members of the seven representative families of large bowel strains in AGORA is also shown. B-E) t-SNE analysis based on individual strain metabolite production/carbohydrate utilisation profile (B: Metabolite production, C: Monosaccharides, D: Oligosaccharides, E: Polysaccharides) suggests more conserved metabolic capabilities at the family level in terms of metabolite production profiles. Datapoint sizes correspond to the number of strains with an overlapping location on a plot, and therefore, and identical metabolic profile. Only strains that were inferred to utilise/produce one or more molecules in a group are included.

Interestingly, despite this variation across AGORA strains, our results suggest that the fermentation/SCFA metabolite production profiles are relatively conserved among members of prevalent gut families, as displayed in Figure 2B where discrete clusters with members of the same family can be observed. In comparison, carbohydrate utilisation profiles show less clustering by family (Figure 2C-E).

Overall, these results suggest a decoupling between functional capabilities and microbial phylogeny in terms of carbohydrate trophic networks. This further highlights the limitations in attempting to design DRC related interventions, or predict intervention outcomes, based on gut microbial composition assessed at higher taxonomic levels.

### Design of an inulin-degrading butyrate-producing community

Applying the functional data that we have generated, we aimed to provide grounding principles to rationally design a microbial purpose-based community to increase metabolite production from a selected DRC. We propose communities at the centre of such interventions should be a) feasible – with a realistic number of members; b) resilient – members should be capable of covering others auxotrophies through cross-feeding during changing nutritional conditions in the gut; and c) function-guided – members should actively contribute to the target metabolite production process, while avoiding detrimental outputs (e.g. hydrogen sulphide production). We define feasible communities as those with 4 up to approximately 30 members, according to existing defined microbial communities in the context of human gut microbiome interventions (66). Based on these characteristics, we aimed to design a minimal community of gut commensals that can enable the production of butyrate from the degradation of inulin (Figure 3A). Inulin is a commonly used and broadly studied polysaccharide (62, 67, 68), while butyrate has been extensively described as a metabolite with a positive impact to gut health (63, 69).

**Figure 3.**
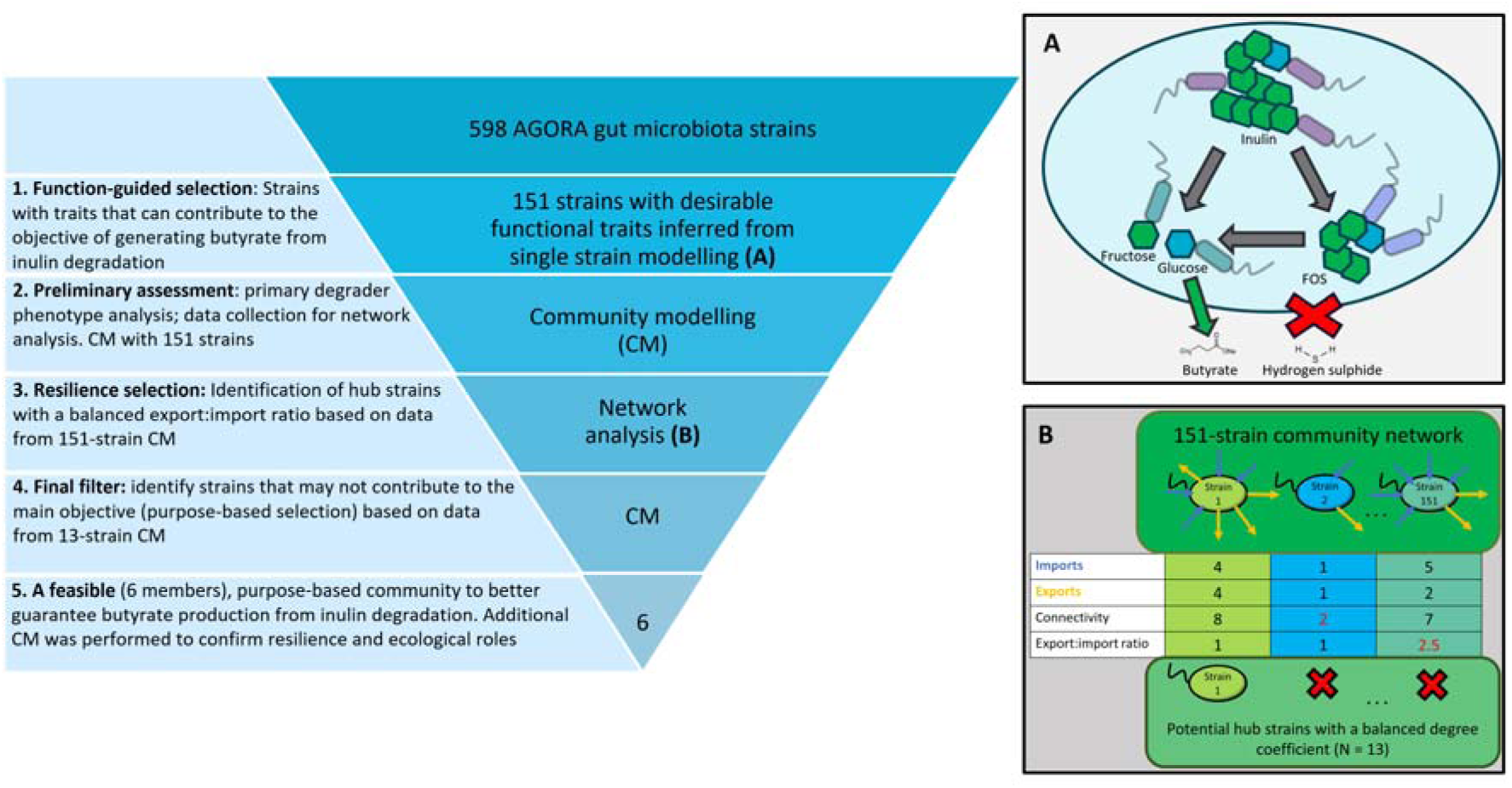
Steps for the design of an inulin-degrading, butyrate producing microbial community. We rely on reverse ecology-derived functional data (step 1 and diagram A), microbial community modelling (steps 2 and 4) and network analysis (step 3 and diagram B) to identify ideal candidates for the implementation of a rational intervention.

We start the consortia design process by using single strain carbohydrate utilisation data generated in the analyses above, identifying every large bowel strain capable of participating in the inulin to butyrate trophic network (Figure 3 step 1 and diagram A; function-guided selection). Based on the metabolic reactions AGORA GEMs can perform, inulin is broken into 29 units of fructose and one unit of glucose or into 25 units of fructose and one unit of kestopentaose, a fructose-oligosaccharide (FOS), which when degraded produces four units of fructose and one unit of glucose. Therefore, we initially selected strains that were identified as inulin and kestopentaose degraders. Additionally, strains capable of generating butyrate from the utilisation of glucose and/or fructose were included. This resulted in an initial selection of 151 strains (Table 1). In doing so, a total of seven different phenotypes were identified, from specialists (inulin or FOS degraders, butyrate producers) to generalists (those that can contribute at every level of the trophic network).

**Table 1.** Predicted functional roles in members of a function-guided 151-strain community. Functional characterisation of AGORA GEMs (single-strain modelling) revealed that 151 strains presented desirable traits to implement this hypothetical intervention, which are grouped by phenotype. Conversely, community modelling of this group of strains identified that not all members presented the traits predicted during single-strain modelling (Retained phenotype: number of strains that retained the predicted function during community modelling). Additionally, specific strains across every relevant phenotype displayed adverse functional traits during community modelling: butyrate consumption and hydrogen sulphide (H_2_S) production.

|  | Single strain modelling | Community modelling |  |  |
| --- | --- | --- | --- | --- |
| Phenotype groups | Total number of strains | Retained phenotype | Butyrate consumers | H<sub>2</sub>S producers |
| Butyrate producers (from glucose and/or fructose) | 60 | 42 | 6 | 11 |
| FOS degraders | 24 | 9 | 0 | 4 |
| Inulin degraders | 33 | 33 | 2 | 8 |
| Inulin-FOS degraders | 16 | 16 | 1 | 5 |
| FOS degraders & butyrate producers | 8 | 4 | 1 | 3 |
| Inulin degraders & butyrate producers | 6 | 6 | 0 | 1 |
| Inulin-FOS degraders & butyrate producers (generalists) | 4 | 3 | 0 | 0 |

To continue with the community refinement process we modelled the community resulting from the 151 selected strains (Figure 3, step 2, Supplementary file 2.7), utilising MICOM, a Python package for the modelling of GEMs as microbial communities (46). The media utilised for modelling was our universal growth media (cfUDM) supplemented with inulin. While predictions showed that every strain grew in this context, results revealed that 10 strains consumed butyrate and that 32 produced hydrogen sulphide (H_2_S, Table 1), a metabolite associated with detrimental impacts on gut health (11, 70). Furthermore, traits predicted in single strain simulations did not always translate to the community setting.

We then performed a flux network analysis, based on the nutritional exchanges predicted in the modelling of the 151 identified strains, to identify those more likely to contribute to a resilient community (Supplementary file 2.8). We calculated total in- and out-degree edges (shared metabolites imported or exported) within the network to identify hub strains, those that are highly interconnected and likely to contribute to community resilience in the tested nutritional context (cfUDM + inulin) (71). We additionally identified strains with a balanced export:import ratio (exports/imports ≈ 1) to prevent the addition of ‘*cheaters*’, strains that take advantage of nutrients available in the media but do not reciprocate accordingly (high imports/low exports). This is supported by the literature, which reports that cheating may contribute to community instability (72). A total of 13 strains were identified to fulfil our network analysis criteria, which were subsequently modelled as a community to assess their ecological roles (Figure 3, step 4; final filter). Community modelling showed that two potential butyrate producers were not predicted to do so in this context, and five strains consumed butyrate and/or produced H_2_S (Supplementary file 2.9). Exclusion of these strains resulted in a purpose-based, six-strain community (Figure 3, step 5).

The resultant community consisted of a primary degrader of inulin (*Prevotella intermedia* 17), four secondary degraders that can produce butyrate (*Clostridium sordellii* ATCC 9714, *Clostridium* sp. SY8519, *Ruminococcaceae* bacterium D16 and *Peptostreptococcus anaerobius* DSM 2949) and a generalist (*Roseburia intestinalis* L1-82). Modelling of our 6-strain purpose-based community (Supplementary file 2.10) confirmed the ecological roles of each strain while also predicting a butyrate production rate (mmol/gDW/h) 30.2% higher than what was predicted for the initial 151-strain selection. A higher inulin to butyrate conversion efficiency was also reported, as the 6-strain community produced 20.2 mmol of butyrate per mmol of inulin degraded (Supplementary file 2.10), while the 151-strain community produced 11.1 mmol of butyrate per mmol of inulin degraded (Supplementary file 2.7).

### Testing resilience in a minimal, purpose-based community

To test the resilience in the predicted purpose-based community, we modelled it in media deprived of proteinogenic amino acids (Figure 4, Supplementary file 2.11). Predictions reported that every strain could grow, supported by complementary prototrophies. Figure 4A shows that *Peptostreptococcus anaerobius* DSM 2949 plays a central role in this nutritional environment as the sole contributor of five amino acids in the restricted media. While every predicted butyrate producer adopts a butyrate consuming phenotype in this extreme scenario, butyrate is still produced albeit at a lower conversion efficiency (6.2 mmol of butyrate produced per mmol of inulin consumed) compared to the amino acid replete media.

**Figure 4.**
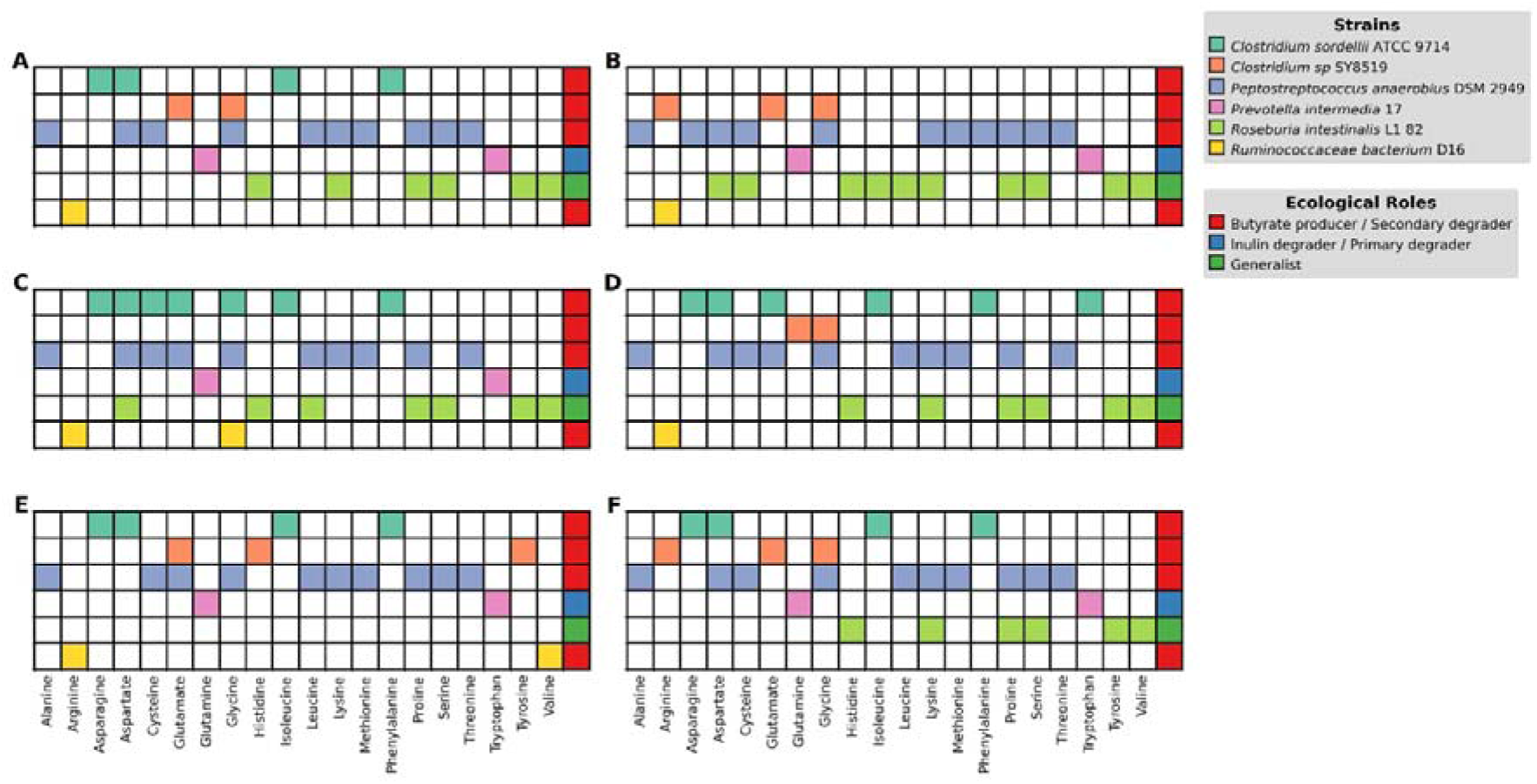
Community resilience in amino acid limitation. A: Modelling of our final inulin-to-butyrate 6-strain community in amino acid restricted media shows that members can cover auxotrophies for the rest of the community while also presenting a degree of functional redundancy. During a knock-one-out analysis (B to F) we identified that the community is resilient to the absence of individual member strains, other than Peptostreptococcus anaerobius DSM 2949 (knockout not shown). The knocked-out strain is shown in grey in panels B to F. Metabolic fluxes for each knock out condition are shown in Supplementary file 3

We additionally tested network resilience by adopting a knock-one-out approach to model the community in the absence of each strain. Importantly, no strain was essential under standard conditions (cfUDM + inulin, Supplementary file 2.12). The overall community also showed resilience to missing one of five of its members under conditions of proteinogenic amino acid restriction (Figure 4B-F, Supplementary file 2.13). Yet, supporting the role of *Peptostreptococcus anaerobius* DSM 2949, it was shown that none of the other five strains were able to grow in amino acid restricted media in the absence of this strain. Together, these results highlight how modelling can inform the design of a resilient community, but also to infer vulnerabilities within the community.

To further support the rationale behind our network analysis selection criteria, we designed a community with a similar structure, however using six strains that did not fit our criteria; four butyrate producers (*Anaerofustis stercorihominis* DSM 17244, *Odoribacter splanchnicus* DSM 20712, *Alcaligenes faecalis* subsp. *Faecalis* NCIB 8687 and *Butyricimonas synergistica* DSM 23225) with the lowest export/import score among the butyrate producing ecological role in our sample. Meanwhile, the selected inulin degrader (*Prevotella intermedia* ATCC 25611) and a generalist (*Roseburia inulinivorans* DSM 16841) also did not meet our requirements regarding reciprocation within a network, and these examples were selected due to their close phylogenetic relationship with the two primary degraders in the original 6-strain community (Figure 3B, poor metabolic contribution to the community). Community modelling in standard media showed that only two strains produce butyrate in this context (one butyrate producer and the community generalist) while one adopts a butyrate consuming phenotype (Supplementary file 2.14). Community butyrate production rate was predicted at a percent change of 60.6% lower than our original 6-strain purpose-based community, while only 12.9 mmol of butyrate was produced per mmol of inulin degraded. In testing the resilience of this network to amino acid limitation, we found that the community had no resilience for methionine restriction, as the amino acid was inferred to be essential for five of the strains. The sixth additional strain (*Alcaligenes faecalis* subsp. *Faecalis* NCIB 8687) whilst not a methionine prototroph, requires alanine from external sources while also relying on primary degraders for access to simpler carbohydrates (Supplementary file 2.15 and 2.16). These findings suggest that the criteria we have devised to identify key members within a complex community can effectively guide the design process for resilient, minimal communities.

### A purpose-based community predicted to improve butyrate production in MCMMs

To infer the potential benefits of the identified purpose-based community in a more realistic setting, we seeded it in 20 predefined species- and 10 genus-level GEMs communities (microbial community-scale metabolic models, MCMMs), mapped from previously characterised human gut microbiome profiles (31). These MCMMs were shown to accurately predict personalised SCFA production when modelled in standard European diet (31) and to positively respond to inulin supplementation. Here, we modelled the impact of seeding these MCMMs with our purpose-based community, using the standard European diet supplemented with 10 mmol of inulin, and maximum butyrate production was predicted (Supplementary file 4).

Predicted maximum butyrate when the MCMMs were supplemented with inulin and the purpose-based community (inulin + module intervention) was found to be higher in most of the predicted MCMMs, as compared to inulin supplementation alone (inulin intervention). The Wilcoxon signed-rank test (Supplementary file 2.17) revealed that this difference was statistically significant (*p* < 0.001, median: 2 mmol/gDW/h, Figure 5A). However, such difference was minimal (<0.1 mmol/gDW/h) in eight samples, which were determined non-responders to the inulin + module intervention. Notably, these were the highest overall butyrate producing MCMMs in the inulin intervention (Figure 5A). Similarly, responders that presented a comparative lower butyrate production in the inulin only intervention displayed the largest increase in butyrate production in the inulin + module intervention. We found that the average number of inulin degraders in native MCMMs (before seeding with the purpose-based community) was lower in responders than non-responders (11.3 versus 12.3 for species-level MCMMs and 5.2 versus 4.6 in genus-level MCMMs).

**Figure 5.**
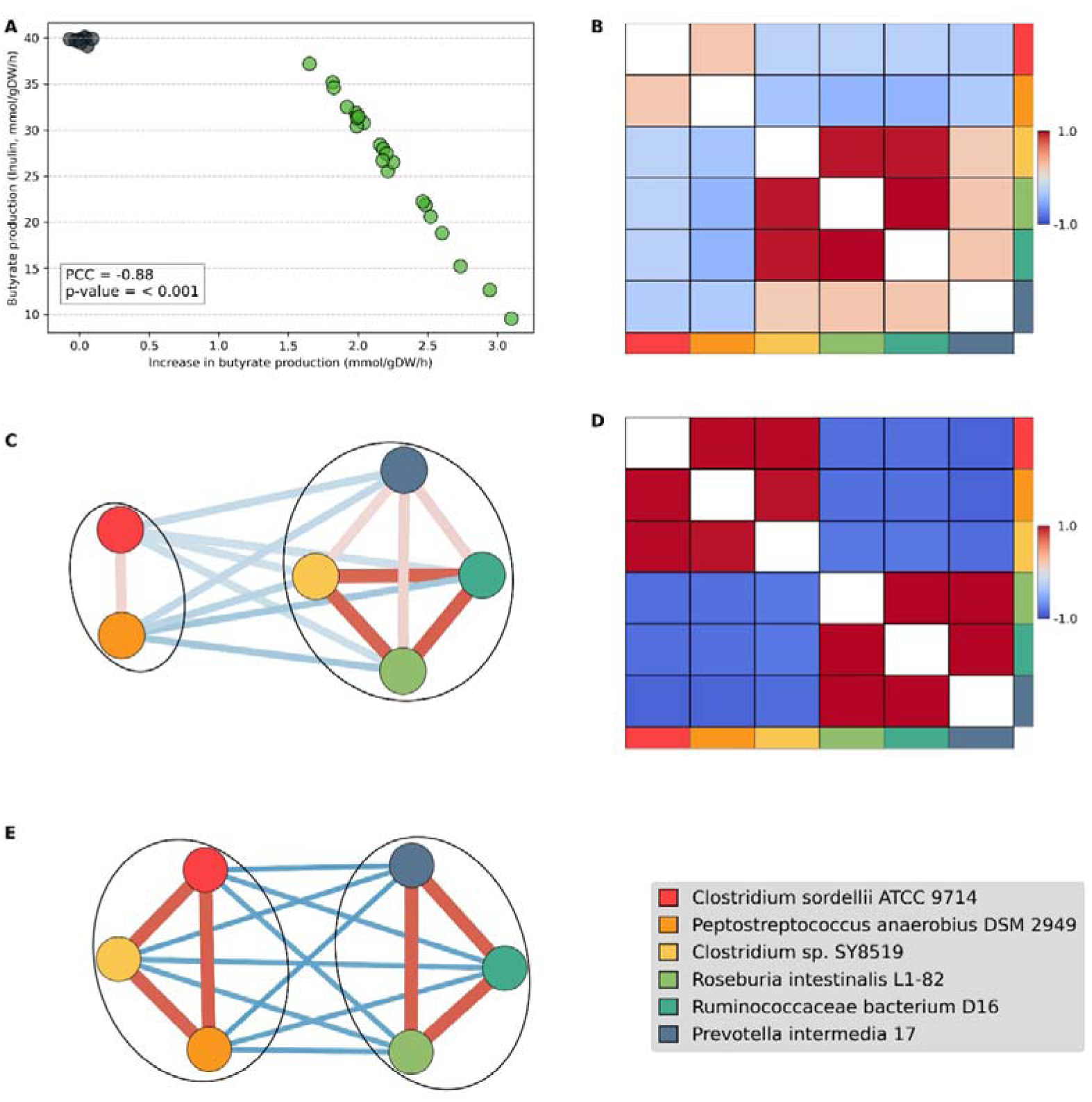
Predicted butyrate and correlative relationship of members of the purpose-based community in modelled MCMMs. A) Scatterplot showing the relationship between maximum butyrate production during the Inulin intervention alone (European diet supplemented with inulin (10 mmol/gDW/h)) versus the predicted increase in maximum butyrate production during the Inulin + Module intervention (purpose-based community at 5% relative abundance) for each MCMMs. Non-responders (butyrate production increment < 0.1 mmol/g/gDW/h) are shown in grey and responders are shown in green. B) Correlation heatmap between members of the purpose-based community in responders. C) Correlation network between members of the purpose-based community in responders. D) Correlation heatmap between members of the purpose-based community in non-responders. E) Correlation network between members of the purpose-based community in non-responders. Positive correlations are shown in red and negative correlations are shown in blue in both heatmaps and networks.

The biomass of three members of the purpose-based community (*Ruminococcaceae bacterium* D16 (butyrate producer), *Clostridium* sp. SY8519 (butyrate producer) and *R. intestinalis* L1-82 (generalist)) was strongly correlated in the MCMMs of responders (PCC > 0.9) while being weakly correlated with the biomass of *P. intermedia* 17 (PCC: 0.21–0.25, Figure 5B-C). The similarities in growth patterns between such strains suggest module-like interdependences. Meanwhile, co-abundances in the non-responding MCMMs suggest that the strains from the purpose-based community form instead two sub-modules, with *Clostridium* sp. SY8519 correlated with *C. sordellii* ATCC 9714 and *P. anaerobius* DSM 2949 (Figure 5D-E). Notably, MCMM fluxes predict that none of the strains from our minimal community produce hydrogen sulphide in any of the samples we modelled.

To better understand the correlative relationship of *Clostridium* sp. SY8519 based on intervention success, we inspected the fluxes from the corresponding GEM in responder and non-responder MCMMs (Supplementary file 4). We found that *Clostridium* sp. SY8519 is a glutamine consumer in responder MCMMs. In contrast, *Clostridium* sp. SY8519 is predicted to export the amino acid in seven of the eight non-responders. Additionally, butyrate production by *Clostridium* sp. SY8519 in non-responder MCMMs is reduced by a factor of 4.34 mmol/h (on average, corrected for strain biomass) compared to responders.

## Discussion

Here, we leverage genome-scale modelling to characterise 816 strains, with potential to occur in the human gut, in terms of their capabilities to utilise a representative group of carbohydrates and the metabolites that can result from this process. Our analysis highlights that DRC utilisation in the gut is an uncommon phenomenon, and a capability that is species, or in some cases, strain-specific. This is coherent, as complex polysaccharide utilisation often requires coordinated synthesis of a range of carbohydrate-active enzymes (73–75), demanding high resource and energy investment (76). This capability may stem from a long-standing evolutionary relationship with the host and its diet as polysaccharide utilisation seems to be a highly specialised trait that provides crucial leverage to select commensals over other organisms (77–79). Yet DRCs are considered key carbon and energy sources for microbes in the larger bowel (26). This implies that DRC utilisers act as energy gatekeepers for the microbial community in the large intestine. Additionally, our findings suggest that metabolite production profiles are highly varied across different strains, as most of the identified metabolite production profiles were shared by four or fewer strains. Hence, community metabolic outputs will be sensitive to both availability of DRCs and microbiome composition, at the species/strain level. While an opportunity to modulate the human gut microbiome exists, this inherent complexity complicates intervention response.

Current technologies allow for the design of microbial communities, which might represent more effective alternatives to native microbiome modulation through fibre supplementation or the use of conventional probiotics. This process can be informed by GEMs modelling. As a proof of principle, we inferred functional traits from AGORA GEMs, and by combining community modelling and network analysis, devised a minimal, resilient microbial community capable of butyrate production from inulin degradation. Our study finds that butyrate production, a key metabolite for colonocyte health (64, 65), is a comparatively uncommon capability among human gut strains represented in AGORA. This further underscores the need to support fibre supplementation with a rationally designed consortia including butyrate producers, that can better guarantee the delivery of intended metabolic outputs.

The framework we propose for the design of purpose-based communities is replicable, traceable and, together with the carbohydrate utilisation dataset we have generated as part of this study, it can be applied for alternative metabolic objectives, and informed by iterative experimental validation. A similar approach could aim to explore different media conditions or strain combinations that additionally minimise the production of metabolites with potential negative impacts. For example, excess succinate, which has been linked to local inflammation when produced to high levels (80, 81). Alternative nutritional dimensions, such as cofactor requirements, should also be accounted for in the design of resilient microbial communities (42). Ultimately, experimental validation of our findings is needed. However, isolating and culturing human gut microbiome commensals remains challenging. Community design efforts focused on previously cultured strains, which also incorporate modelling approaches such as we have described, will result in a more targeted and directed exploration of the combinatorial space.

The resulting purpose-based community described here is based on a hierarchical structure, where its members have distinct metabolic roles, aiming to contribute to a shared functional output and to community resilience. This is based on the findings of our carbohydrate utilisation assessment of individual AGORA strains, that highlight the determinant role primary degraders have initiating DRC breakdown. In addition, strains can adopt different metabolic strategies, that lead to the production and export of alternative metabolites, even when carbon sources remain unchanged. Therefore, our proposed community encompasses two inulin degraders and four butyrate producers. Members of the purpose-based community are also predicted to be capable of metabolic burden distribution in amino acid restricted conditions. Substantial literature underscores functional redundancy, metabolic synergy and distribution of metabolic burden as key characteristics in stable microbial communities (82–85). Yet, the community we design is shown to rely on *P. anaerobius* DSM 2949 to cover a number of amino acid auxotrophies, which is not necessarily a desirable trait from a resilience perspective; therefore, the addition of an extra member to distribute this burden might be beneficial. Nevertheless, it is likely that increasing the number of total community members exponentially increases its complexity and, hence, should be carefully considered.

Seeding the minimal community into 30 MCMMs, validated for their accuracy in predicting SCFA production (31), and supplemented with inulin, led to a significant increase in predicted butyrate production (2 mmol/gDW/h) in approximately 75% of cases. There were also 8 non-responding MCMMs identified. Our results suggest that communities that would benefit the most from this type of intervention (inulin + module) are the ones that present lower butyrate production levels at baseline (inulin only intervention). This effect has been reported before across microbiomes and in ecological theory (86–88). We also identify compositional differences between responding and non-responding MCMMs. While we do not identify a correlation between intervention response and the biomass of inulin degraders in the purpose-based community (*R. intestinalis* L1-82 and *P. intermedia* 17), we do find that non-responders have a higher number of inulin degraders on average, than responders. It is possible that a limited number of niches for primary degraders exist and that the introduction of additional competitors negates any beneficial effects. A preliminary analysis of the 30 MCMMs showed them to positively respond to inulin supplementation alone. This aligns with findings by Quinn-Bohmann *et al.,* where only 16 MCMMs are reported as non-responders to a high-fibre diet from a sample of 3,129 (31).

The purpose-based community displayed cohesive dynamics when modelled in isolation. Yet, when modelled as part of pre-designed MCMMs, it was predicted to display two configurations composed of sub-modules. Additionally, the correlative relationship of *Clostridium* sp. SY8519 was found to vary with intervention response. Flux analysis suggests that glutamine cross-feeding may partially explain such configurations and intervention responsiveness, as glutamine production in *Clostridium* sp. SY8519 was also associated with reduced butyrate production by this species in non-responders. Interestingly, glutamine supplementation has been shown to impact gut microbiome composition in humans and animal models (89, 90). Overall, these findings highlight the challenges of introducing a defined community into the highly diverse human gut microbiome. Additional mechanisms that better explain this response and the decoupling of *C. sordellii* ATCC 9714 and *P. anaerobius* DSM 2949 remain to be investigated in future studies. Yet, these predicted dynamics suggest that a lower number of secondary degraders can lead to more concise outcomes. The results also highlight the potential importance of designing individualised interventions. The ability to reach such a detailed level of system inspection highlights the value of GEMs-based approaches for hypothesis generation and the study of complex biological systems.

Nevertheless, GEMs still face limitations. For example, it is possible that the limited count of butyrate producers in AGORA is explained or influenced by challenges related to gaps in genome annotation of transporter gene polymorphisms and lack of experimental data, resulting in GEMs that do not encode the corresponding pathways (91, 92). Additionally, our study characterises carbohydrate utilisation capabilities in AGORA strains qualitatively. While a quantitative assessment of such capabilities would have provided greater insights limited experimental data prevents reliable mapping of enzymatic rates. MICOM partially addresses this challenge through the application of its cooperative_tradeoff method, which significantly reduces the simulation space (46). Despite these limitations, genome-scale modelling is a rapidly growing technology with an ever-expanding repertoire of GEMs and efforts to develop GEMs-based frameworks that better capture cellular and community dynamics are ongoing (35, 93); for example, the integration of meta-transcriptomics and GEMs community modelling (35) or the application of AI to improve model accuracy (91, 93). Importantly, predicted relative differences and percentage changes can provide meaningful insights of the studied system (94, 95). Recently, a quantitative metabolite exchange scoring system was proposed to identify exchanged metabolites that are central in a community of GEMs (11). Meanwhile, the conceptual framework we propose here focuses on the identification of strains/GEMs, in a given network, that can better guarantee the establishment of resilient minimal communities. Proof of concept GEMs-based interventions of the human gut microbiome have been hypothesised (31), however, to the best of our knowledge, this is the first GEMs pipeline for informing community-based interventions and their design.

## Conclusion

Through applying genome-scale metabolic modelling we have provided a system-level overview of the metabolic capabilities of human gut microbes in the carbohydrate nutritional dimension, and applied this as functional building-blocks for intervention design. We introduce a comprehensive community design framework that encompasses strain selection, resilience inference, and expands previous work for the prediction of intervention response. In doing so, we have set the stage for subsequent studies to refine and expand upon such rationale. Frameworks such as we have presented here, capable of enabling mechanistic insights into human gut microbiome dynamics, will continue to contribute to our understanding of its role and applications in human health.

## Declarations

## Data availability

All data generated or analysed during this study are included in this published article and its supplementary information files. The datasets generated and/or analysed during the current study are available in the purpose-based community design repository, https://bitbucket.csiro.au/scm/~mol131/purpose-based-community-design.git

## Authors’ contributions

J.M., M.R., E.S. conceived of the presented idea. J.M. reviewed the literature, generated and analysed GEMs data and performed networks analyses. J.M., M.R., D.M., and E.S. were major contributors in writing the manuscript with input from all authors. All authors read and approved the final manuscript.

## Acknowledgements

Not applicable

## Supplementary material

### Supplementary file 1

Description: Supplementary tables and figures.

Format: Word file (.docx)

### Supplementary file 2

Description: Single strain and community modelling data results and data analysis.

Format: Excel file (.xlsx)

### Supplementary file 3

Description: Modelling fluxes from knock-one-out analysis in the 6-strain resilient community.

Format: Excel file (.xlsx)

### Supplementary file 4

Description: Pre- and post-intervention MCMM modelling fluxes.

Format: Zipped folder (.zip)

### Supplementary file 5

Description: Phylogeny tree file for large bowel strains in AGORA.

Format: Tree file (.phy)

