## Supplementary file 1 for "Genome scale metabolic modelling of human gut microbes to inform rational community design"

### Supplementary Tables

#### Table S1 Composition of Universally Defined Media (UDM), engineered to guarantee growth of every GEM in AGORA, and carbohydrate-free UDM (cf-UDM), to allow determination of carbohydrate utilisation capabilities in AGORA GEMs. Nutrients that were removed from UDM to form sf-UDM are shown in red.

| **Nutrient category** | **Nutrients** |
| --- | --- |
| Carbohydrates | Glucose, fructose, galactose, mannose, ribose, lactose, fucose, xylose, arabinose, acetyl glucosamine, glucosamine, chitobiose, inulin, maltose, sucrose, D-galacturonate |
| Amino acids | D-alanine, L-alanine, asparagine, aspartate, arginine, cysteine, cystine, glutamine, glycine, glutamate, histidine, isoleucine, leucine, lysine, D-methionine, L-methionine, L-methionine sulfoxide, phenylalanine, proline, D-serine, L-serine, threonine, tryptophan, tyrosine, valine, L-carnitine, ornithine |
| Dipeptides | Alanyl-glutamine, carnosine, cysteinylglycine, glycyl-L-asparagine, glycyl-L-glutamine, glycylleucine, glycyl-L-methionine, spermidine, glycilcysteine, glycyl-L-tyrosine, glycyl-phenylalanine, L-alanyl-L-threonine,  L-methionyl-L-alanine, 'L-alanyl-L-leucine', glycylproline, L-alanyl-L-aspartate, L-alanylglycine, alanyl-glutamate, glycyl-L-aspartate, glycyl-L-glutamate |
| Fatty acids | Stearic acid, myristic acid, dodecanoic acid, oleic acid |
| Bile acids | Chenodeoxycholic acid, glycocholic acid, taurocholic acid |
| Cations | Calcium, cadmium, mercury, magnesium, sodium, ammonia, potassium, hydrogen ion, nitrogen |
| Anions | Chloride ion, phosphate, sulphate, sulphite, hydrogen sulphide, hydrogen, thiosulphate, nitrite, nitrate |
| Metals | Copper, ferrum2, ferrum3, cobalt, manganese, nickel, zinc |
| Main cofactors | Biotin, cobalamin I, cobalamin II, adenosylcobalamin, folic acid, tetrahydrofolic acid, menaquinone 7, menaquinone 8, demethymenaquinone, ubiquinone 8, nicotinic acid, niacinamide, nicotinamide ribotide, pantothenic acid, riboflavin, reduced riboflavin, pyridoxine, pyridoxal, pyridoxamine, thiamine, thiamine monophosphate |
| Secondary cofactors | Heme, siroheme, thymidine, cytosine, uracil, adenosine, adenine, guanine, deoxyadenosine, deoxyguanosine, guanosine, guanosine triphosphate, methylthioadenosine, adenosine monophosphate, S-adenosylmethionine, deoxyadenosine triphosphate, 5-thymidylic acid, hypoxanthine, cytidine, inosine, xanthine, deoxycytidine, uridine, deoxyinosine, cytidine monophosphate, xanthosine |
| Other | 4-aminobenzoate, glutathione diaminoheptanedioate, dephospho-CoA, 1,2-diacyl-sn-glycerol, methyl-oxovaleric acid, chorismite, 4-hydroxybenzoic acid, oxidized glutathione, putrescine, indole, lanosterin, choline sulfate, ketobutyric acid, glycolaldehyde, trimethylamine, acetic acid, formic acid, gamma-butyrobetaine, acetoacetic acid, coenzyme A, ethanolamine, tetrathionate,  dehydro-deoxy-gluconate, carbon dioxide, allantoin, cholesterol, formaldehyde, water, phenylpyruvic acid, urea, L-lactic acid, citric acid, malic acid, acetylmannosamine, glycerol |

#### Table S2 Carbon sources selected to assess metabolic capabilities in AGORA strains. 66 molecules were selected in total and divided into four categories based on structure complexity.

| **Category** | **Compounds** |
| --- | --- |
| Polysaccharides | Amylose, amylopectin, arabinan, larch arabinogalactan, arabinoxylan, β-glucans, cellulose, dextran, carob galactomannan, glycogen, homogalacturonan, inulin, levan, lichenin, laminarin, α-mannan, pectin, pectic galactan, pullulan, potato rhamnogalacturonan I, wine rhamnogalacturonan II, starch, xylan, xyloglucan. |
| Oligosaccharides | Arabinotriose, cellobiose, N,N-diacetylchitobiose, kestopentaose, kestotetraose, α-lactose, D-maltose, maltohexaose, mannotriose (β-1,4), melibiose, raffinose, starchyose, sucrose, trehalose |
| Monosaccharides | N-acetylgalactosamine, N-acetyl-D-glucosamine, N-acetylneuraminic acid, D-arabinose, L-arabinose, deoxyribose, D-fructose, L-fucose, glucosamine, D-glucose, L-xylose, D-mannose, D-ribose, L-rhamnose, salicin, D-xylose |
| Metabolites | Acetic acid, acetaldehyde, butyrate, ethanol, formic acid, fumaric acid, lactic acid, malic acid, propionic acid, pyruvic acid, succinic acid |

#### Table S3. Classification of strains in AGORA based on their relationship to the host and typical topological (microbiome) location. *The group of 29 AGORA strains more typically associated with the human skin microbiome was excluded from further assessment.

| Category | Definition | Number of GEMs |
| --- | --- | --- |
| Upper GIT | Strains prevalent in the oral cavity, stomach and first portion of the small intestine | 66 |
| Large bowel | Strains commonly identified as part of the human gut microbiome | 598 |
| Pathogens | Strains identified as either pathogens or pathobionts | 123 |
| Skin* | Strains more typically associated with the human skin microbiome | 29 |

### Supplementary Figures

#### Figure S1 Nutrient utilisation capabilities

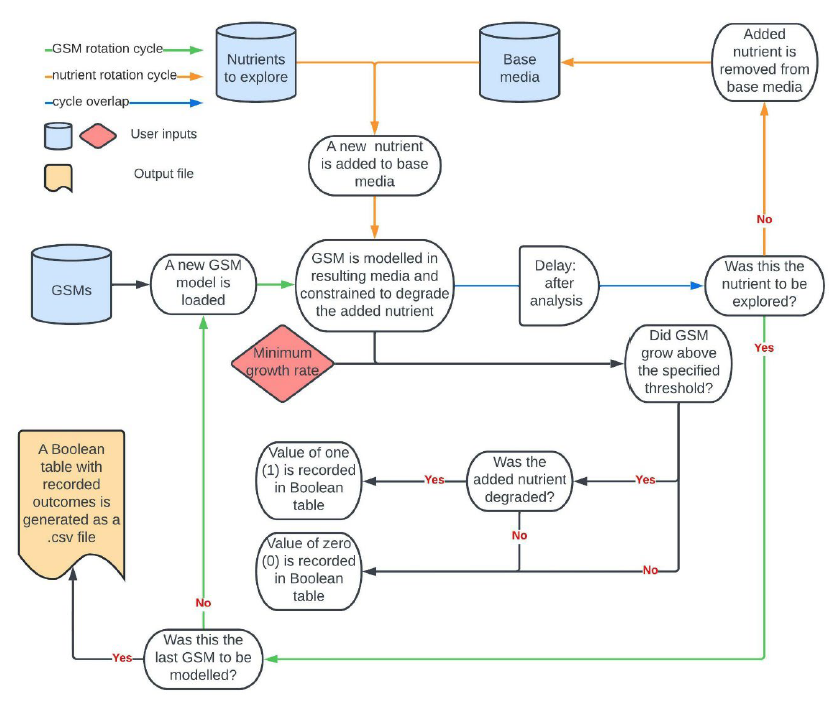

#### Nutrient utilisation capabilities (technical description):

1.	**Inputs:**

a.	**GEMs Directory (Input A):** A folder containing the GEMs to be analyzed.

b.	**Base Media Ingredients (Input B):** A list of exchange reactions representing the base media.

c.	**Test Molecules (Input C):** A list of exchange reactions for molecules not included in the base media (i.e. carbohydrates to be tested).

2.	**Methodology:**

a.	For each GEM in the directory:

i.	For each molecule in Input C:

1.	**Media Preparation:** Create a modified media by adding the molecule to the base media.

2.	**Simulation:** Perform FBA to simulate growth.

3.	**Constraint Application:** Constrain the model (setting the uptake rate of the molecule's exchange reaction to a positive value) to consume the added molecule, if possible.

4.	**Outcome Determination:**

a.	If the model grows above a predefined threshold and consumes the molecule, confirmed by flux inspection, assign a positive outcome (1).

b.	If not, assign a negative outcome (0).

3.	**Data Organization:**

a.	Compile results into a Boolean matrix with strains as rows and test molecules as columns.

#### Figure S2 Metabolite export capabilities following nutrient utilisation

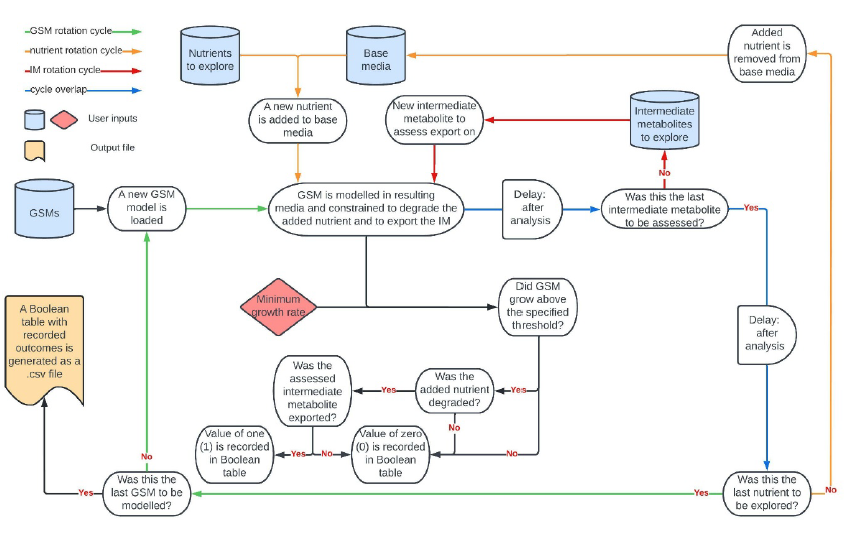

#### Metabolite export capabilities following nutrient utilisation (technical description)

1.	**Inputs:**

a.	**GEMs Directory (Input A):** A folder containing the GEMs to be analyzed.

b.	**Base Media Ingredients (Input B):** A list of exchange reactions representing the base media.

c.	**Test Molecules (Input C):** Molecules to be tested for utilization (i.e. carbohydrates to be tested).

d.	**Target Metabolites (Input D):** List of metabolites to assess for export.

2.	**Methodology:**

a.	For each GEM in the directory:

i.	For each molecule in Input C:

1.	For each metabolite in Input D:

a.	**Media Preparation:** Add the molecule to the base media.

b.	**Nutrient Constraint:** Constrain the model to utilise the molecule.

c.	**Export Constraint:** Constrain the model to produce the metabolite, if metabolically feasible (setting a minimum flux for the metabolite's exchange reaction, if feasible, constraining the model to produce and export such metabolite during simulation).

d.	**Simulation:** Perform FBA to simulate growth.

e.	**Outcome Determination:**

i.	If the model grows above the threshold, consumes the molecule, and exports the metabolite, all confirmed by flux inspection, assign a positive outcome (1).

ii.	If not, assign a negative outcome (0), this includes cases where a phenotype that follows the specified metabolic constraints is unfeasible.

3.	**Data Organization:**

a.	Compile results into a Boolean matrix with strains as rows and combinations of molecules and metabolites as columns.

#### Figure S3 Carbohydrate utilisation experimental design

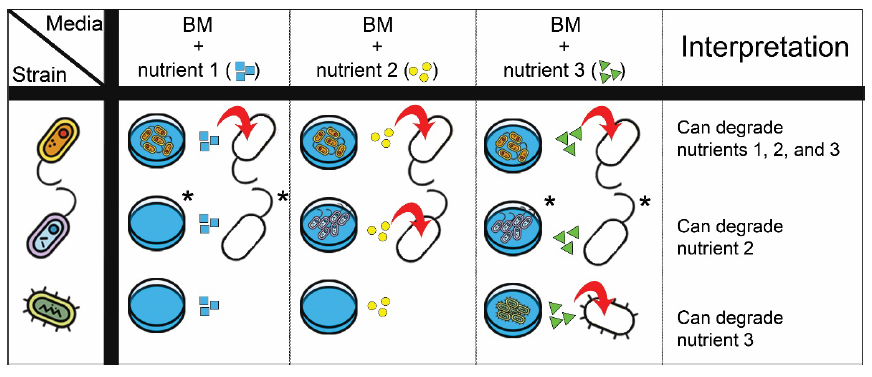

#### Figure S4 Intermediate metabolite export experimental design

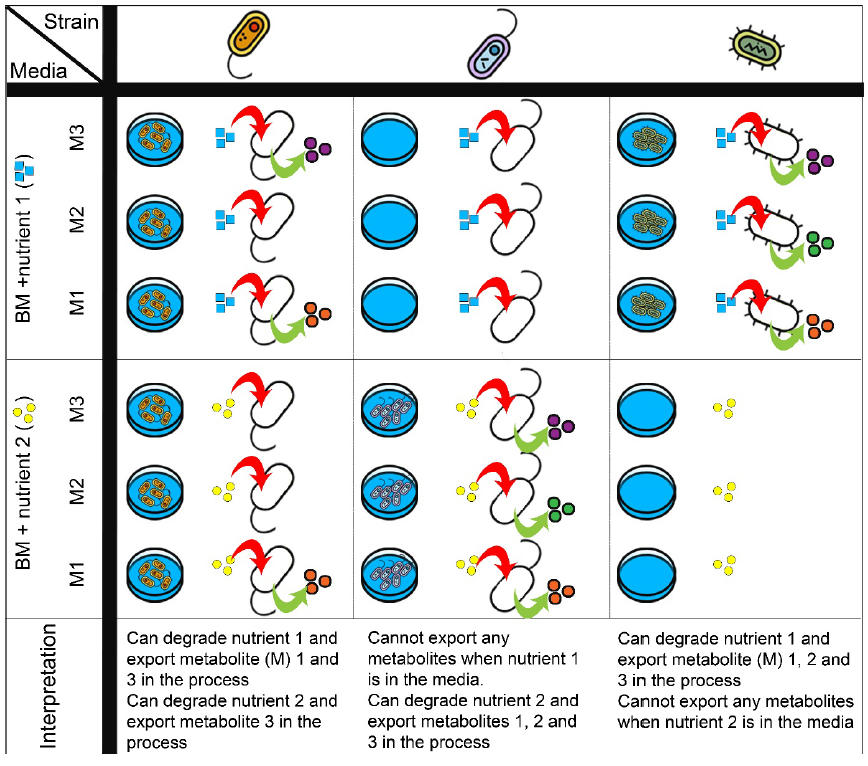

#### Figure S5 Distribution of carbohydrate utilisation capabilities among topo-physiological groups of AGORA strains.

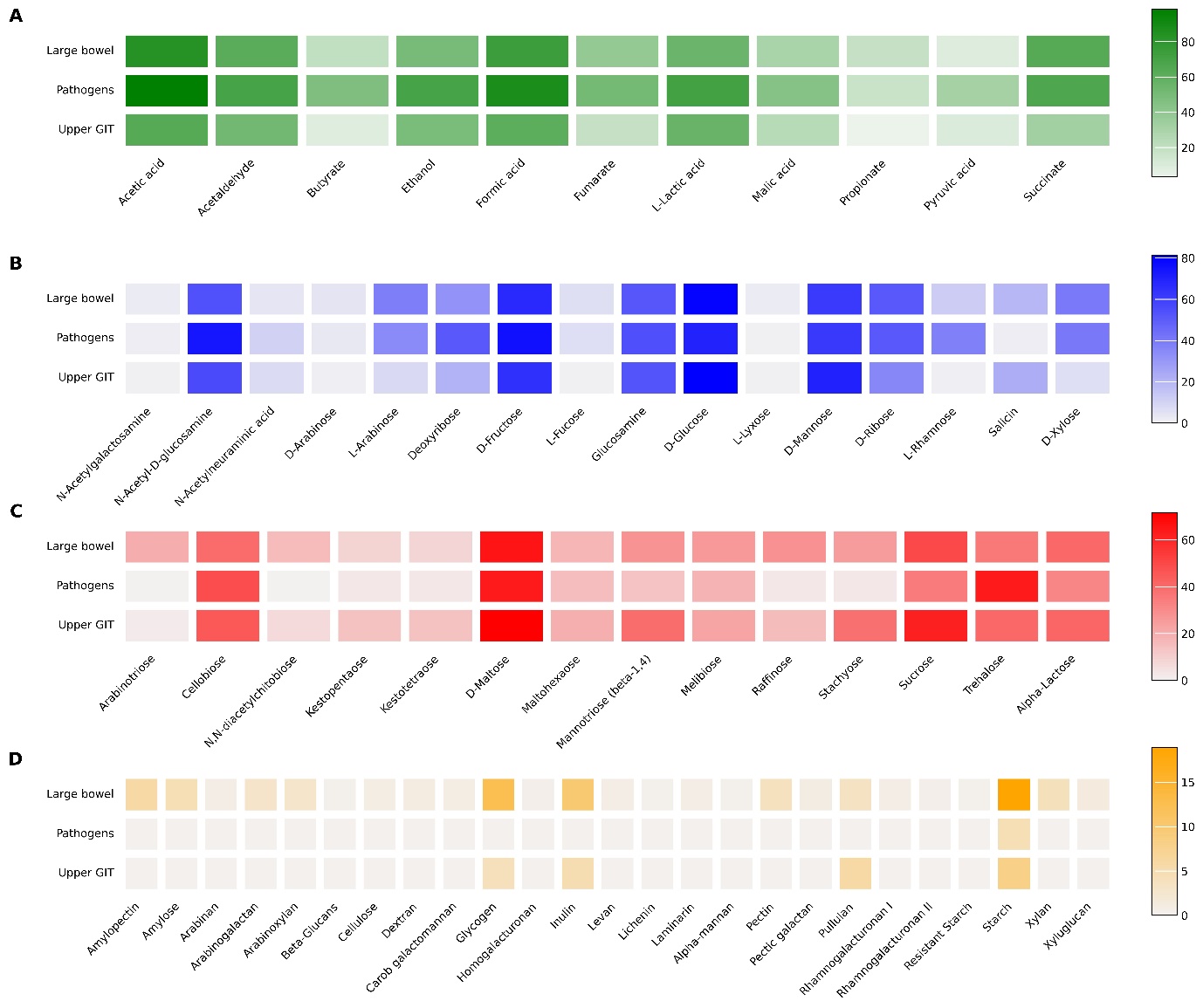

Heatmaps show the percentage of strains from each group inferred to utilise a given nutrient from one of our four structural categories (A: Intermediate metabolites, B: Monosaccharides, C: Oligosaccharides, D: Polysaccharides), enabling the identification of functional traits within each group. Darker hues indicate higher percentages. Analysis of the functional differences between topo-physiological groups, allowed us to establish trends in the carbohydrate nutritional dimension (Figure 1). Pathogens show higher propensity, in terms of percentage of strains, for the utilisation of intermediate metabolites (Figure 1A) as compared to the large bowel or upper GIT groups. Pathogens also show higher propensity than strains in the large bowel group for utilising select monosaccharides including those found in mucin (N-acetyl-D-glucosamine, N-acetylneuraminic acid, fucose) and deoxyribose, (Figure 1B), mostly an intracellular metabolite. However, as nutrient structural complexity increases, members of the Pathogens group demonstrate reduced utilisation capacity. For example, this group of strains shows almost null capabilities to utilise polysaccharides, with the exception of starch (Figure 1D). Conversely, strains in the Upper GIT group display higher affinity for oligosaccharides such as D-maltose, mannotriose, sucrose and lactose (Figure 1C) and for specific monosaccharides such as glucose and mannose, all of which are common components of diet but also absorbed by the host in the small intestine. Finally, Large bowel strains dominate in terms of polysaccharide utilisation over the other groups as well as the monosaccharides that derive from their utilisation. This highlights access to DRC-derived nutrients/energy as a key aspect of niche adaptation in the gut

#### Figure S6 Inferred metabolite production by monosaccharide utilised

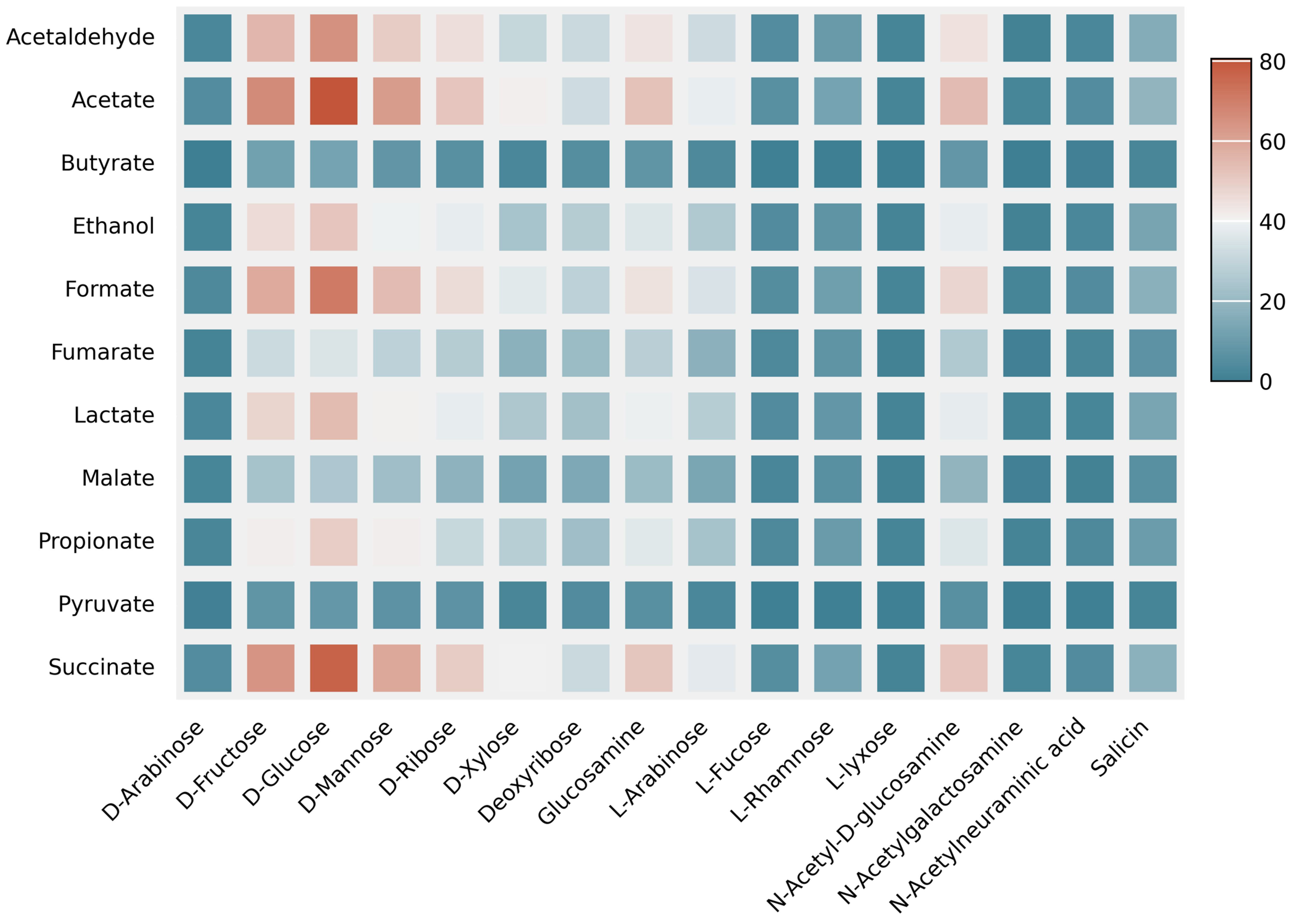

##### *The heatmap shows the percentage of large bowel strains inferred to produce each of the 11 tested metabolites while utilising a specific monosaccharide.*
